# Layer-specific developmentally precise axon targeting of transient suppressed-by-contrast retinal ganglion cells (tSbC RGCs)

**DOI:** 10.1101/2021.11.26.470118

**Authors:** Nai-Wen Tien, Tudor C. Badea, Daniel Kerschensteiner

## Abstract

The mouse retina encodes diverse visual features in the spike trains of more than 40 retinal ganglion cell (RGC) types. Each RGC type innervates a specific subset of the more than 50 retinorecipient brain areas. Our catalog of RGC types and feature representations is nearing completion. Yet, we know little about where specific RGC types send their information. Furthermore, the developmental strategies by which RGC axons choose their targets and pattern their terminal arbors remain obscure. Here we identify a genetic intersection (*Cck-Cre* and *Brn3c^CKOAP^*) that selectively labels transient Suppressed-by-Contrast (tSbC) RGCs, a member of an evolutionarily conserved functionally mysterious RGC subclass. We find that tSbC RGCs selectively innervate the dorsolateral and ventrolateral geniculate nuclei of the thalamus (dLGN and vLGN), the superior colliculus (SC), and the nucleus of the optic tract (NOT). They binocularly innervate dLGN and vLGN but project only contralaterally to SC and NOT. In each target, tSbC RGC axons occupy a specific sublayer, suggesting that they restrict their input to specific circuits. The tSbC RGC axons span the length of the optic tract by birth and remain poised there until they simultaneously innervate their four targets around postnatal day five. The tSbC RGC axons make no errors in choosing their targets and establish mature stratification patterns from the outset. This precision is maintained in the absence of Brn3c. Our results provide the first map of SbC inputs to the brain, revealing a narrow target set, unexpected laminar organization, target-specific binocularity, and developmental precision.

**Significance statement:** In recent years, we have learned a lot about the visual features encoded by retinal ganglion cells (RGCs), the eye’s output neurons. In contrast, we know little about where RGCs send their information and how RGC axons, which carry this information, target specific brain areas during development. Here, we develop an intersectional strategy to label a unique RGC type, the tSbC RGC, and map its projections. We find that tSbC RGC axons are highly selective. They innervate few retinal targets and restrict their arbors to specific sublayers within these targets. The selective tSbC RGC projection patterns develop synchronously and without trial and error, suggesting molecular determinism and coordination.

## Introduction

Vision begins in the retina, which transforms the pixel representations of photoreceptors into feature representations of retinal ganglion cells (RGCs), the eye’s sole output neurons (Gollisch and Meister, 2010). The mouse retina contains more than 40 RGC types, which send different visual information to the brain (Baden et al., 2016; Bae et al., 2018; Rheaume et al., 2018; Tran et al., 2019; Goetz et al., 2021). We have learned a lot about how retinal circuits extract visual features and encode them in specific RGC types’ spike trains (Schwartz, 2021). Yet, we know little about where this information is sent.

In total, mouse RGCs innervate more than 50 brain areas (Morin and Studholme, 2014; Martersteck et al., 2017). However, which brain areas specific RGC types innervate remains, with few exceptions, unknown (Hattar et al., 2006; Huberman et al., 2008, 2009; Kim et al., 2008; Yonehara et al., 2008). Some RGC axons target distinct layers within the dorsolateral geniculate nucleus (dLGN) and superior colliculus (SC), the two largest retinorecipient areas of the mouse brain (Reese, 1988; Huberman et al., 2009; Kim et al., 2010; Hong et al., 2011; Kerschensteiner and Guido, 2017). Our maps of RGC axons in dLGN and SC are incomplete, and the retinal input organization of many other targets is unknown. Finally, a subset of RGCs (9/40+ types) innervate ipsi- and contralateral brain areas to support binocular vision (Dräger and Olsen, 1980; Johnson et al., 2021). Whether these RGC types innervate all or only a subset of their targets binocularly is unknown.

Retinal projections to the brain serve as a model for studying the development of long-range neural projections and supported the discovery of conserved growth programs and molecular cues that guide RGC axons towards their targets (Erskine and Herrera, 2007; Moore and Goldberg, 2011; Mason and Slavi, 2020; Williams et al., 2020). Yet, how RGC axons invade specific targets and organize their arbors within, and whether this process involves trial-and-error or proceeds orderly, is unknown (Kim et al., 2010; Osterhout et al., 2011, 2014, 2015; Su et al., 2021). Individual RGCs innervate multiple targets through axon collaterals to ensure complete visual field representation in each retinorecipient area (Fernandez et al., 2016). Whether collaterals innervate targets simultaneously or sequentially is unclear (Osterhout et al., 2014). Brn3 POU domain transcription factors are part of the gene regulatory network that controls RGC specification and terminal identity (Mu and Klein, 2004; Badea et al., 2009). Brn3a, Brn3b, and Brn3c are expressed in distinct but overlapping sets of RGCs (Xiang et al., 1995; Badea and Nathans, 2011; Parmhans et al., 2018, 2021). RGC axon targeting is severely affected in Brn3b KO mice, but the contributions of Brn3a and Brn3c to this process remain unknown (Badea et al., 2009, 2012).

Most RGCs increase their firing rates to light increments (i.e., positive contrast), light decrements (i.e., negative contrast), or both. However, in 1967, Rodieck and Levick discovered RGCs in cats and rabbits, respectively, that fire at high rates in the absence of stimuli and are silenced by positive and negative contrast steps (Levick, 1967; Rodieck, 1967). Suppressed-by-Contrast (SbC) RGCs have since been identified in non-human primates and mice, indicating that they are a conserved output of the mammalian retina (de Monasterio, 1978; Jacoby et al., 2015; Tien et al., 2015). The contributions of SbC signals to vision remain mysterious, in part because the projection patterns of SbC RGCs are unknown (Masland and Martin, 2007).

Here, we discover a genetic intersection that selectively labels transient SbC (tSbC) RGCs in mice. We use this intersection to map the tSbC RGC projections to the brain and study their development.

## Materials and Methods

### Experimental animals

We used the following mouse lines, alone or in combination: *Cck-Cre* (Taniguchi et al., 2011), *Brn3c^CKOAP^* (Badea et al., 2012), and *Ai9* (Madisen et al., 2010). All mice were crossed onto a *C57Bl6/J* background for at least five generations prior to inclusion in this study.

### Tissue preparation and histochemistry

Animals were given an overdose of anesthesia and perfused transcardiacally with ice cold PBS followed by 4% paraformaldehyde (Sigma). Fixed brain and retina vibratome slices and fixed retinal whole mounts, were washed twice for 20 minutes in PBS at room temperature. Before alkaline phosphatase (AP) staining, brain sections and retinas in PBS were heated in a water bath at 65-70 °C for 90 minutes to inactivate endogenous AP. AP staining was then developed in the heat-inactivated tissue in AP buffer (0.1M Tris, 0.1M NaCl, 50 mM MgCl2, pH to 9.5) with 3.4 μL/mL of NBT/BCIP overnight at room temperature with gentle agitation (Badea et al., 2003). After staining, the tissue was washed three times for 20 minutes in PBS with 0.1% Tween 20 and fixed overnight in PBS with 4% PFA in 4 °C. To improve imaging, the tissue was dehydrated through an ethanol series (50%, 75%, 85%, 95%, and 100% [100 proof] for 20 minutes, then 100% [200 proof] overnight) and cleared with 2:1 Benzyl benzoate (BB)/Benzyl alcohol (BA). The tissue was then mounted in 2:1 BB/BA between glass slides and coverslips. before the NBT/BCIP precipitate were dissolved in BB/BA. We used a rabbit polyclonal anti-Brn3c antibody (Xiang et al., 1995) to stain *Cck-Cre Ai9* retinas.

### Imaging and analysis

We brightfield imaged the cleared tissue with 4X and 10X objectives on an Olympus BX51 microscope. We used an Olympus FV1000 laser-scanning confocal microscope with a 20X objective to analyze Brn3-staining in *Cck-Cre Ai9* mice.

Density recovery profiles (DRPs) of labeled RGCs were calculated following the definitions of Rodieck (1991) using scripts written in MATLAB. RGC dendrite and axon stratification profiles were analyzed in Fiji (Schindelin et al., 2012). To determine whether labeled RGCs are regionally enriched or distributed evenly across the retina, we measured their density in four quadrants of each retina and defined an asymmetry index as:

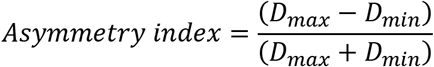

where, *D_max_* is the density of cells in the most populated quadrant and *D_min_* is the density of cells in the least populated quadrant.

## Results

### An intersectional genetic strategy selectively labels tSbC RGC

We previously characterized tSbC RGCs in *Cck-Cre* transgenic mice injected with adeno-associated viruses (AAVs) or crossed to a fluorescent reporter strain (*Ai9,* tdTomato) (Tien et al., 2015). The bistratified dendrites of tSbC RGCs target synaptic laminae outside the inner plexiform layer’s (IPL’s) ChAT bands (i.e., the plexus of ON and OFF starburst amacrine cells) and are connected by numerous ascending and descending processes (Tien et al., 2015). An RGC type matching this morphological description was reported in *Brn3c^CKOAP/+^* mice crossed to a ubiquitously expressed sparsely induced Cre line (*R26CreERT* mice) (Badea and Nathans, 2011). Both *Cck-Cre Ai9* and *R26CreERT Brn3c^CKOAP/+^* label multiple RGC types. To examine their overlap, we first stained *Cck-Cre Ai9* mice for Brn3c. This co-labeled a set uniformly sized RGC somata (Fig. 1*A*). We next crossed *Cck-Cre* and *Brn3c^CKOAP/+^* mice. In mature *Cck-Cre Brn3c^CKOAP/+^* mice (postnatal day 30, P30), a sparse population of RGCs (172 ± 7 cells / mm^2^, n = 10 retinas) expressed alkaline phosphatase (Fig. 1*B*). The density recovery profiles (DRPs) of the labeled RGCs showed pronounced exclusion zones (Fig 1*C*), a hallmark of cell-type-specific retinal mosaics (Rodieck, 1991). Cells were distributed evenly across the retina (Fig. 1*D*). In vibratome slices, the labeled RGCs were consistently bistratified with dendrite arbors outside the expected ChAT band positions (Fig. 1*E,F*) (Bae et al., 2018). Thus, the intersection of *Cck* and *Brn3c* in *Cck-Cre Brn3c^CKOAP/+^* mice selectively labels tSbC RGCs.

**Figure 1.**
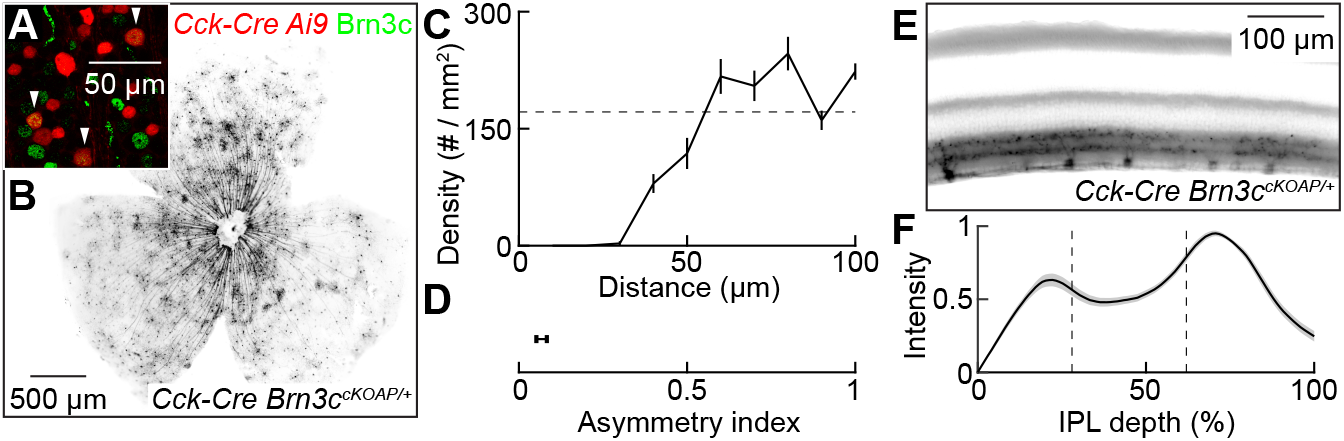
*Cck-Cre Brn3c^CKOAP/+^* mice selectively label tSbC RGCs. ***A,*** Excerpt of a flat-mounted P30 *Cck-Cre Ai9* (red) retina stained for Brn3 (green). ***B,*** Retinal whole mount from a P30 *Cck-Cre Brn3c^CKOAP/+^* mouse. ***C,*** Density recovery profile (DRP) of RGCs labeled in P30 *Cck-Cre Brn3c^CKOAP/+^* mice. Line indicates the mean and errorbars the SEM (n = 10 retinas). ***D,*** Asymmetry index for the distribution of labeled RGCs in the four retinal quadrants (see Materials and Methods, n = 5 retinas). ***E,*** Vibratome slices through a 30 *Cck-Cre Brn3c^CKOAP/+^* retina. ***F,*** Labeling profile across the IPL depth (i.e., RGC dendritic stratification profile) in P30 *Cck-Cre Brn3c^CKOAP/+^* mice. Line indicates the mean and shaded areas the SEM (n = 10 retinas).

### tSbC RGCs innervate four brain areas in a layer-specific manner

To map the projections of tSbC RGCs, we cut coronal sections of *Cck-Cre Brn3c^CKOAP/+^* mouse brains (P30) and stained them for alkaline phosphatase. Neurons in the brain did not express alkaline phosphatase. All brain labeling was eliminated by binocular enucleation (*data not shown),* indicating that it reflects the expression of alkaline phosphatase in tSbC RGC axons.

We comprehensively surveyed the retinorecipient areas of *Cck-Cre Brn3c^CKOAP/+^* mouse brains (P30) and found labeling in only four (of more than 50) (Morin and Studholme, 2014; Martersteck et al., 2017). In the LGN complex, the dLGN and ventrolateral geniculate nucleus (vLGN) were labeled, whereas IGL was clear (Fig. 2*B*). Labeling in dLGN encompassed all but the most medial aspect, and in vLGN, labeling was restricted to the most temporal layer (Fig. 2*B*). In SC, labeling was strongest in the upper layer of the retinorecipient superficial SC (sSC, Fig. 2*C*). Finally, we observed narrowly stratified labeling in the nucleus of the optic tract (NOT, Fig. 2*D*). Thus, tSbC RGC axons innervate a small subset of retinorecipient areas (4/50+) and target specific layers within each of these areas.

**Figure 2.**
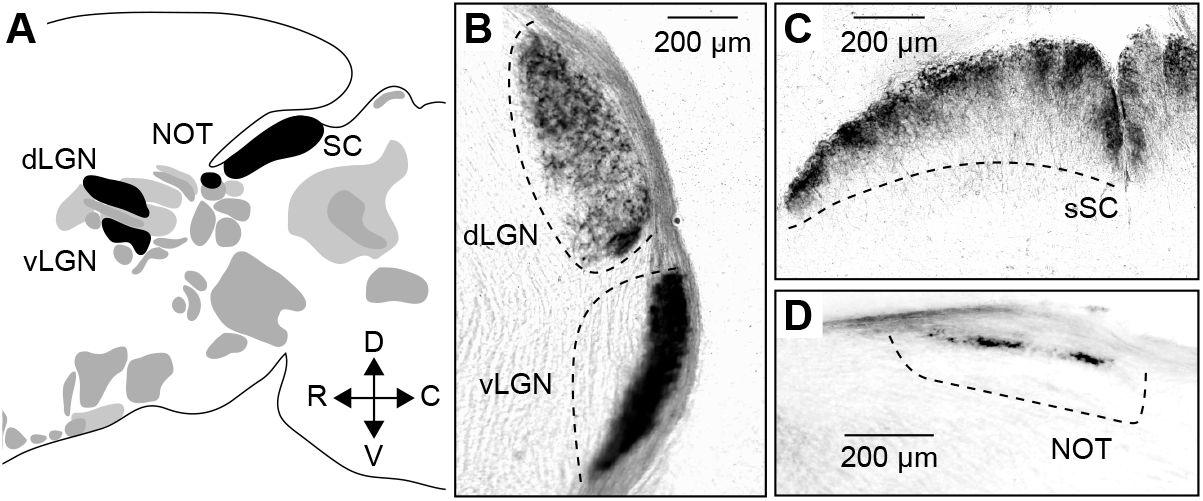
tSbC RGCs innervate four brain areas in a laminar-specific manner. ***A,*** Sagittal schematic view of the brain illustrating retinorecipient brain areas, with those innervated by tSbC RGCs painted black. ***B-D,*** Coronal sections of the LGN complex (***B***), SC (***C***), and NOT (***D***) of a P30 *Cck-Cre Brn3c^CKOAP/+^* mouse showing the laminar-specific axonal targeting of tSbC RGCs.

### tSbC RGCs binocularly innervate a subset of their targets

We recently discovered that only nine of the more than 40 mouse RGC types innervate ipsi- and contralateral brain areas to support binocular vision (Johnson et al., 2021). The remaining types innervate only contralateral targets. tSbC RGCs are part of the ipsilaterally projecting set (Johnson et al., 2021). To test whether tSbC RGCs innervate all their targets binocularly, we examined alkaline phosphatase labeling in the brains of monocularly enucleated *Cck-Cre Brn3c^CKOAP/+^* mice. We enucleated mice at P25 and analyzed RGC projection patterns at P30 to allow for Wallerian degeneration of RGC axons from the removed eye (Knöferle et al., 2010). All four targets received contralateral tSbC RGC input (Fig. 3*A-C*), but only the dLGN and vLGN received input from ipsilateral tSbC RGCs (Fig. 3*A*). Thus, tSbC RGCs binocularly innervate only a subset of their targets, the first example of target-specific RGC binocularity outside the special case of the retinohypothalamic tract (Magnin et al., 1989; Fernandez et al., 2016).

**Figure 3.**
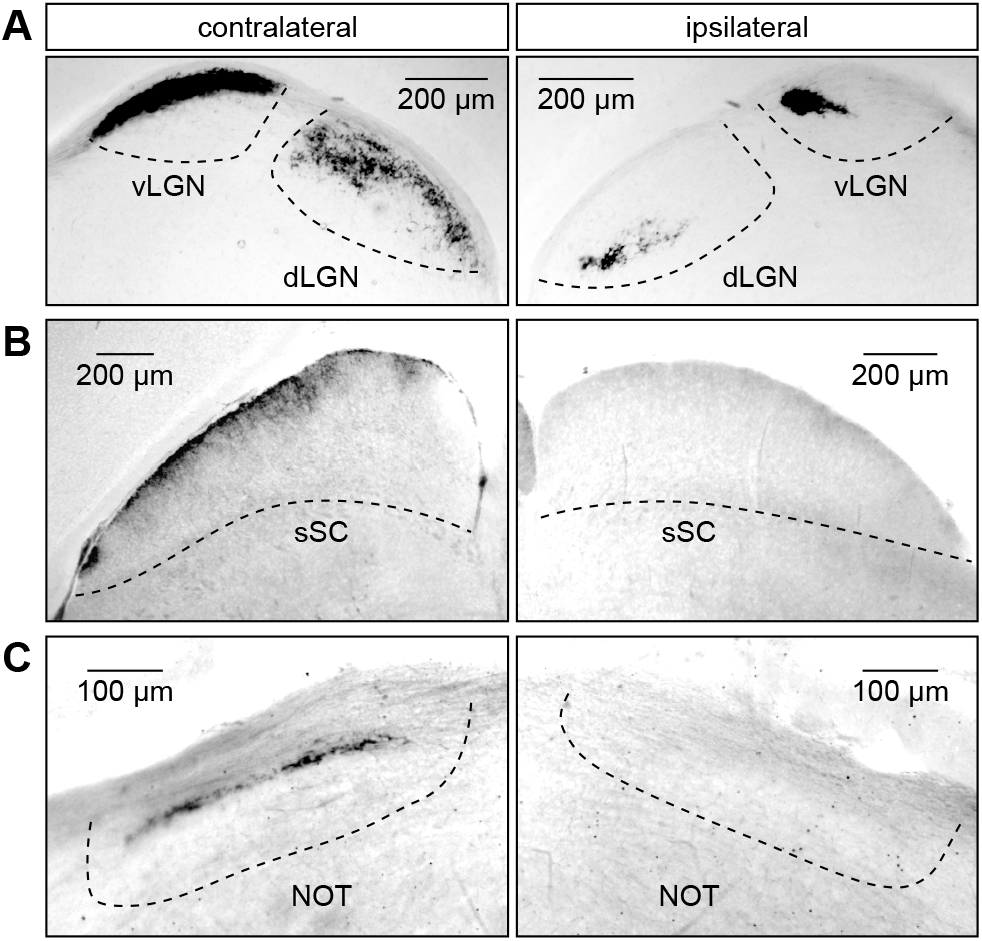
tSbC RGCs binocularly innervate a subset of their targets. ***A-C,*** Coronal sections of contra (left) and ipsilateral (right) LGN (***A***), SC (***B***), and NOT (***C***) five days after enucleation of a P30 *Cck-Cre Brn3c^CKOAP/+^* mouse.

### *Cck-Cre Brn3c^CKOAP/+^* mice label tSbC RGCs throughout postnatal development

To analyze how tSbC RGCs establish target- and layer-specific axonal projections, we first needed to test whether *Cck-Cre Brn3c^CKOAP/+^* mice selectively label these cells during development. We prepared retinal whole mounts from P0, P5, P10, P15, and P20 mice to address this question. At P0, some photoreceptors, in addition to RGCs, expressed alkaline phosphatase (Fig. 4*A*); at all other ages, labeling was restricted to RGCs (Fig. 4*B-E*). The density of labeled RGCs increased with age (P0: 96 ± 8 cells / mm^2^, P5: 109 ± 12 cells / mm^2^, P10: 130 ± 8 cells / mm^2^, P15: 171 ± 19 cells / mm^2^, P20: 215 ± 15 cells / mm^2^, n = 10 retinas for all ages). Most importantly, the RGCs’ DRPs showed clear exclusion zones at all ages (effective radius, P0: 25.7 ± 1.7 μm, P5: 30.8 ± 1.5 μm, P10: 30.9 ± 1.4 μm, P15: 27.7 ± 1.2 μm, P20: 29.8 ± 1.3 μm), indicating that *Cck-Cre Brn3c^CKOAP/+^* mice selectively label tSbC RGCs throughout postnatal development (Fig. 4*F-J*).

**Figure 4.**
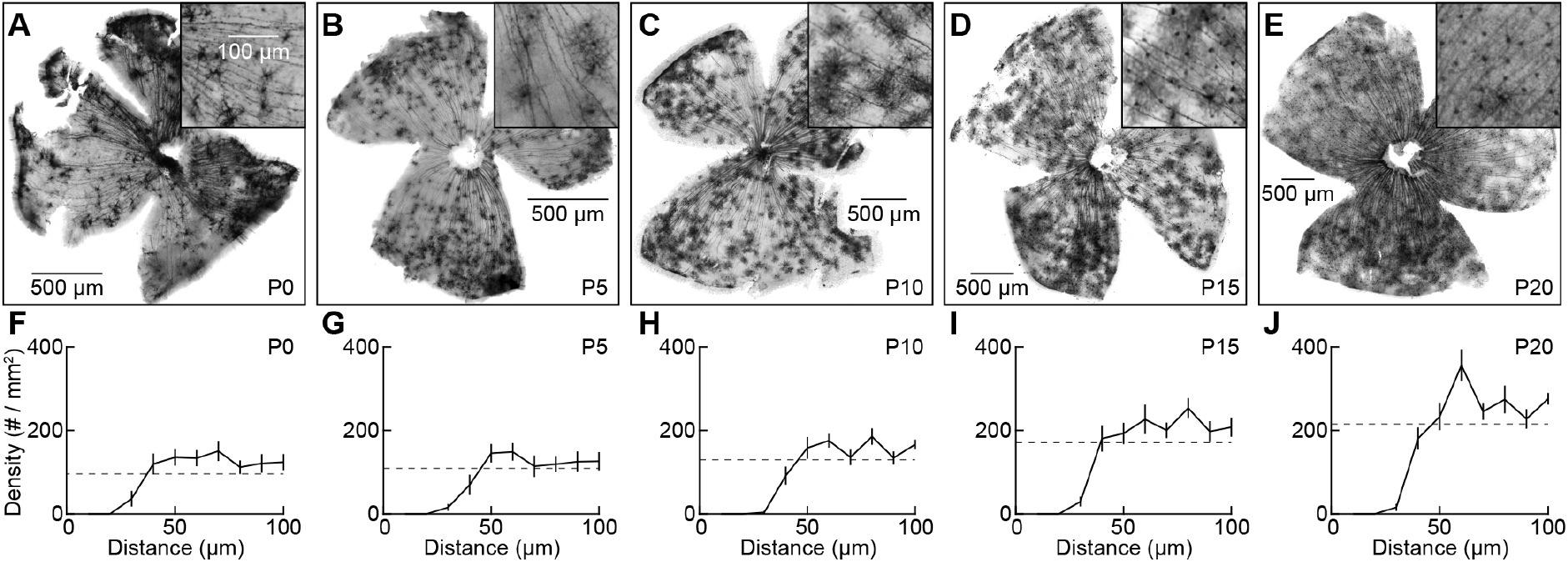
*Cck-Cre Brn3c^CKOAP/+^* mice label tSbC RGCs throughout postnatal development. ***A-E,*** Representative whole mounts of *Cck-Cre Brn3c^CKOAP/+^* retinas at P0 (***A***), P5, (***B***), P10 (***C***), P15 (***D***), and P20 (***E***). Insets show magnified views of smaller regions at mid-eccentricity. ***F-J,*** DRPs of labeled RGCs in *Cck-Cre Brn3c^CKOAP/+^* retinas at P0 (***F***), P5, (***G***), P10 (***H***), P15 (***I***), and P20 (***J***) (n = 10 retinas for each).

### Development of the tSbC RGC projections

We started to study the development of tSbC RGC projections by tracking their innervation of the LGN complex from P0 to P20. At P0, labeling was restricted mainly to the optic tract with few branches or axon terminals visible in the dLGN or vLGN (Fig. 5*A-C*). However, by P5 axonal arborizations occupied all but the medial aspect of dLGN and a lateral band of vLGN (Fig. 5*A-C*). These patterns were maintained to maturity (Fig. 5*A-C*). We observed no aberrant tSbC RGCs axons in the intergeniculate leaflet at any stage of postnatal development.

**Figure 5.**
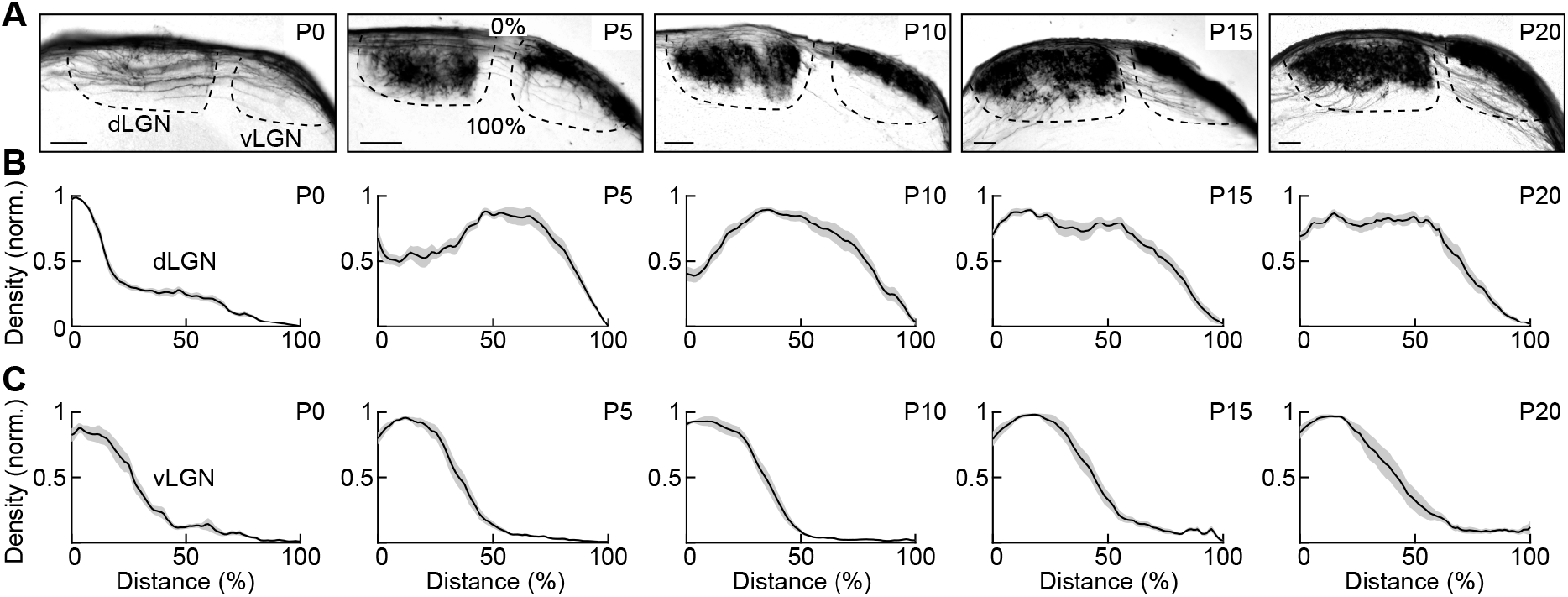
Development of the tSbC RGCs’ LGN innervation. ***A,*** Developmental time-series of coronal sections through the LGN complex of *Cck-Cre Brn3c^CKOAP/+^* mice (P0, P5, P10, P15, P20, from left to right). ***B,C,*** Labeling profiles from the optic tract (0%) to the medial boundaries (100%) of dLGN (***B***) and vLGN (***C***) across development (P0, P5, P10, P15, P20, from left to right). Lines indicate the means and shaded areas the SEMs (n = 3-5 mice at each timepoint).

We next turned to the SC. Like LGN, tSbC RGC axons had few branches in sSC at P0 but established mature arborization patterns by P5 (Fig. 6*A-C*). The tSbC RGC axons occupied the uppermost layers of sSC from P5 to maturity (Fig. 6*A-C*). We found that neurons in the deeper layers SC (dSC) expressed alkaline phosphatase during early postnatal development (P0 and P5). We confirmed that the sSC labeling reflects tSbC RGC axons entering from the optic tract in sagittal sections (Fig. 7).

**Figure 6.**
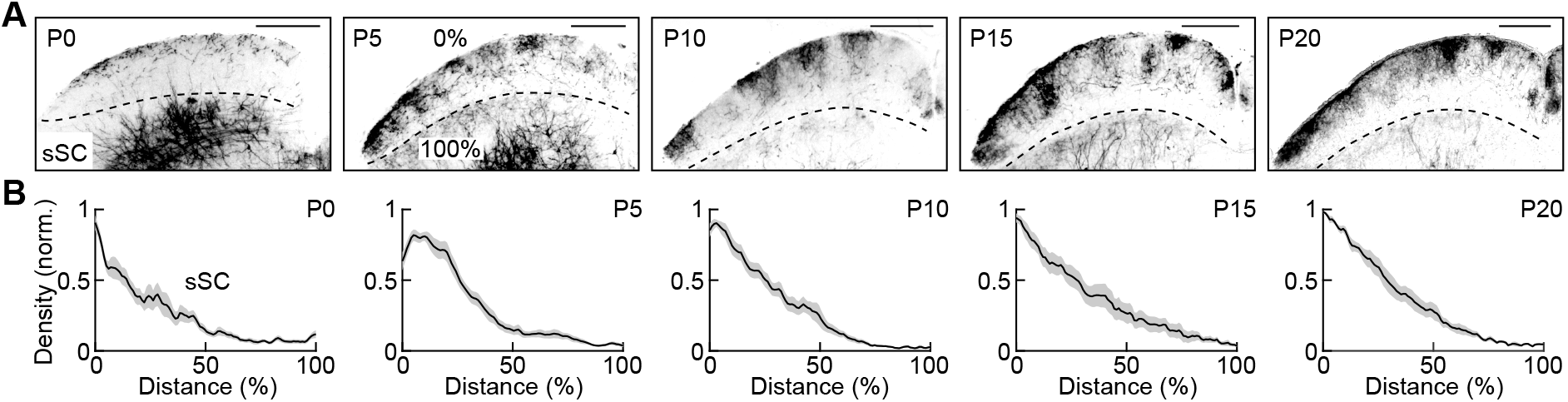
Development of the tSbC RGCs’ SC innervation. ***A,*** Developmental time-series of coronal sections through the SC of *Cck-Cre Brn3c^CKOAP/+^* mice (P0, P5, P10, P15, P20, from left to right). ***B,*** Labeling profiles from the surface of SC (0%) to the caudal margin of sSC (i.e., the caudal margin of the stratum opticum, 100%) across development (P0, P5, P10, P15, P20, from left to right). Lines indicate the means and shaded areas the SEMs (n = 3-5 mice at each timepoint).

**Figure 7.**
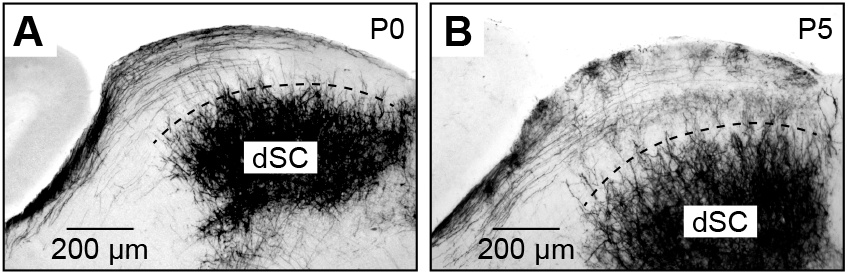
Labeling in *Cck-Cre Brn3c^CKOAP/+^* sSC represents retinal axons. ***A,B,*** Sagittal sections through the SC of P0 and P5 *Cck-Cre Brn3c^CKOAP/+^* mice. At these ages, neurons in dSC transiently express alkaline phosphatase. However, the neurites of dSC neurons are sparse in sSC and most of the sSC labeling reflects axons entering from the optic tract (i.e., tSbC RGC axons).

Finally, although tSbC RGC axons were present in the optic tract at P0, we observed few branches in the NOT (Fig. 8*A-C*). By P5, tSbC RGC axon terminals targeted a narrow band in NOT and maintained this position to maturity (Fig. 8*A-C*). At no point in our developmental time series did we find tSbC RGCs axons in other pretectal areas or any retinorecipient targets other than dLGN, vLGN, SC, and SC NOT. Thus, tSbC RGC axons choose their targets without developmental errors, invade them simultaneously between P0 and P5, and form laminar arborizations patterns that are precise from the outset.

**Figure 8.**
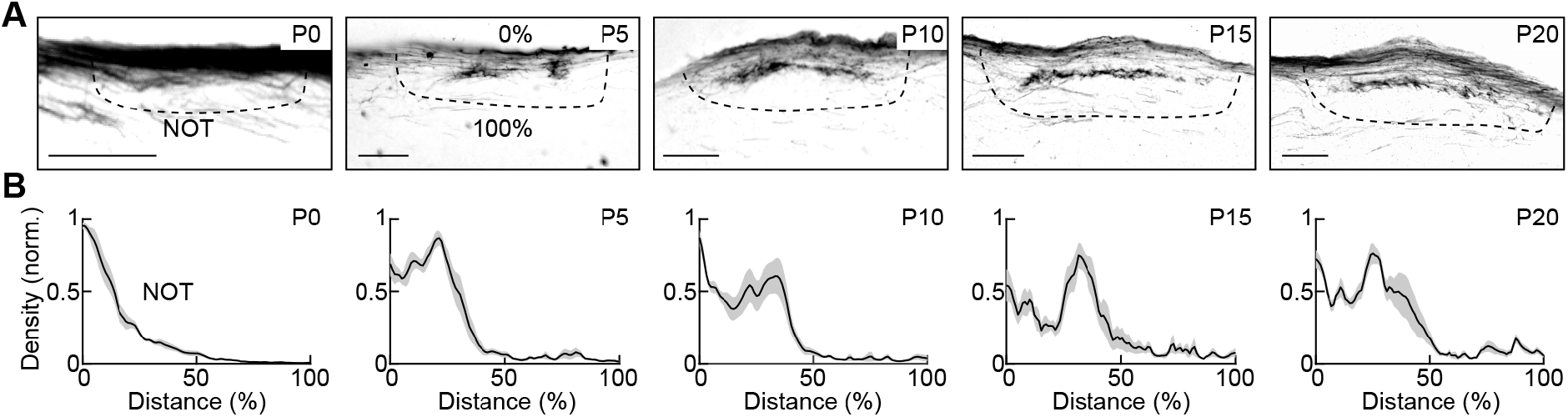
Development of the tSbC RGCs’ NOT innervation. ***A,*** Developmental time-series of coronal sections through the NOT of *Cck-Cre Brn3c^CKOAP/+^* mice (P0, P5, P10, P15, P20, from left to right). ***B,*** Labeling profiles from the optic tract (0%) to the caudo-medial margin of NOT (100%) across development (P0, P5, P10, P15, P20, from left to right). Lines indicate the means and shaded areas the SEMs (n = 3-5 mice at each timepoint).

### tSbC RGC mosaics and axonal projections develop independently of Brn3

Brn3 transcription factors contribute to RGC differentiation and terminal identity (Mu and Klein, 2004; Badea et al., 2009). Brn3a, Brn3b, and Brn3c are each expressed in multiple RGC types, and their contributions to RGC differentiation and axon targeting have not been studied with cell-type resolution. We generated *Cck-Cre Brn3c^CKOAP/CKOAP^* mice to explore the influence of Brn3c on tSbC RGCs. We observed no differences in the density of tSbC RGCs (Fig. 9*A,B*, *Cck-Cre Brn3c^CKOAP/CKOAP^:* 205 ± 5 cells / mm^2^, n = 5 retinas, *Cck-Cre Brn3c^CKOAP/+^:* 215 ± 15 cells / mm^2^, n = 10, p = 1 by Mann-Whitney U test), their mosaic distributions (Fig. 9*A,B*, exclusion zones, *Cck-Cre Brn3c^CKOAP/CKOAP^:* 31.7 ± 1.2 μm, n = 5 retinas, *Cck-Cre Brn3c^CKOAP/+^:* 29.8 ± 1.3 μm, n = 10, p = 0.21 by Mann-Whitney U test), or axonal targeting (Fig. 9*C-H*), between P20 *Cck-Cre Brn3c^CKOAP/CKOAP^* and *Cck-Cre Brn3c^CKOAP/+^* mice. We do not know when in development, Brn3c is removed in these mice but know that Cre-dependent alkaline phosphatase expression is detectable at least from P0 onwards. Thus, the postnatal development of tSbC RGCs and their axonal projections is independent of Brn3c.

**Figure 9.**
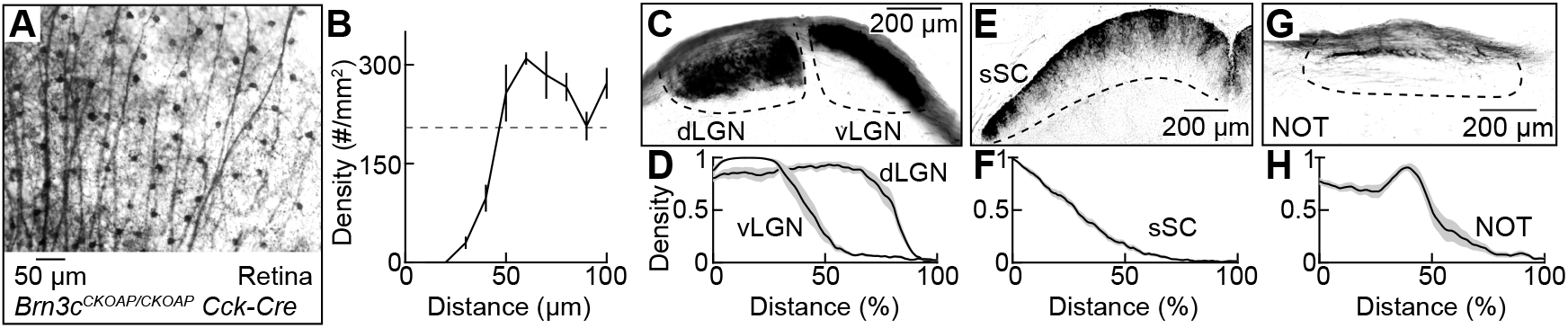
tSbC RGC mosaics and axonal projections develop independently of Brn3c. ***A,*** Excerpt of a P30 *Cck-Cre Brn3c^CKOAP/CKOAP^* retina showing alkaline phosphatase expression in the tSbC RGC mosaic. ***B,*** DRP of RGCs labeled in P30 *Cck-Cre Brn3c^CKOAP/CKOAP^* mice (n = 5 retinas). ***C,E,G,*** Coronal sections of the LGN complex (***C***), SC (***E***), and NOT (***G***). ***D,F,H,*** Labeling profiles from the optic tract (0%) to the medial boundaries of dLGN and vLGN (***D***, 100%), the surface of SC (0%) to the caudal margin of sSC (***F**,* 100%), and the optic tract (0%) to the caudo-medial margin of NOT (***H***, 100%). Lines indicate the means and shaded areas the SEMs (n = 3 mice).

## Discussion

Here, we discover a transgenic intersection (*Cck-Cre* and *Brn3c^CKOAP^*) that marks a single RGC type in the mouse retina, the tSbC RGC. Immunohistochemistry (Fig. 1) and single-cell RNA sequencing support the notion that tSbC RGCs co-express *Cck* and *Brn3c* (Tran et al., 2019; Goetz et al., 2021). We use *Cck-Cre Brn3c^CKOAP^* mice to map tSbC RGC projections and study their development.

The tSbC RGCs selectively innervate four of the more than 50 retinorecipient brain areas: dLGN, vLGN, SC, and NOT (Fig. 2). The dLGN and SC are the main image-forming retinal targets (Morin and Studholme, 2014; Martersteck et al., 2017). Both pass information to visual cortex (among other places), dLGN directly to primary visual cortex (V1), and SC via the lateral posterior nucleus of the thalamus (LP) to postrhinal cortex (POR) (Kerschensteiner and Guido, 2017; Beltramo and Scanziani, 2019; Bennett et al., 2019). SbC responses have been recorded in dLGN, SC, and V1 (Niell and Stryker, 2008; Piscopo et al., 2013; Ito et al., 2017), suggesting that some dLGN and SC neurons receive exclusive or predominant input from tSbC RGCs to relay their signals to visual cortex and other second-order targets. In addition, high-resolution imaging of RGC axon responses suggests that SbC inputs converge with conventional retinal signals onto some dLGN neurons (Liang et al., 2018). The combination of SbC and conventional signals could serve as a contrast gain control mechanism in which SbC signals boost responses to low-contrast stimuli while avoiding response saturation to high-contrast stimuli.

The NOT is part of the accessory optic system, which controls gaze-stabilizing eye movements via the optokinetic reflex arc (Simpson, 1984). The NOT, in particular, mediates ipsiversive horizontal eye movements (Kato et al., 1986; Yakushin et al., 2000; Macé et al., 2018). NOT neuron light responses have, to our knowledge, not been characterized in mice. In rabbits, they are unfailingly direction-selective (Collewijn, 1975). NOT is strongly innervated by ON direction-selective RGCs in rabbits and in mice (Pu and Amthor, 1990; Dhande et al., 2013). Thus, it seems likely that tSbC RGCs converge with ON direction-selective RGCs onto NOT neurons, possibly for contrast gain control. It seems less likely that tSbC RGC signals are maintained as a separate information stream in this target. Why tSbC RGCs do not innervate medial terminal nucleus (MTN) - the nucleus of the accessory optic system mediating gaze-stabilizing vertical eye movements - is unclear.

The vLGN has recently been shown to mediate influences of light on mood and learning (Huang et al., 2019, 2020). The vLGN also regulates defensive responses to visual threats (Fratzl et al., 2021; Salay and Huberman, 2021), but this regulation appears to be independent of its light responses and retinal input (Fratzl et al., 2021). The vLGN is subdivided into a retinorecipient external (vLGNe) and a non-retinorecipient internal (vLGNi) layer. Interestingly, a recent study identified transcriptionally distinct cell types arranged in four sublaminae within vLGNe (Sabbagh et al., 2020). Our labeling distribution suggests that tSbC RGCs provide input to the outer two layers of vLGNe (Figs. 2 and 5).

The laminar targeting of axons in the other retinorecipient areas suggests that tSbC RGCs provide input to specific circuits in these targets as well. It will be interesting to see if and where tSbC RGCs axons overlap with axons of other SbC-RGC types (Jacoby and Schwartz, 2018; Goetz et al., 2021). Our discovery of a genetic intersection to target tSbC RGCs should allow us in the future to selectively perturb these cells to clarify their contributions to visual processing and behaviors mediated by the targets identified here.

A previous study found that early-born RGCs’ axons grow along the optic tract prenatally and transiently innervate more targets than they ultimately maintain (Osterhout et al., 2014). In contrast, late-born RGCs’ axons grow along the optic tract postnatally and make few if any targeting errors (Osterhout et al., 2014). We find that tSbC RGC axons span the length of the optic tract at P0 indicating that they grow prenatally. They remain poised there until ~P5 when they simultaneously and without error innervate their four targets P5 (Figs. 5-8), revealing further cell-type-specific diversity in the developmental strategies of retinofugal projections.

Some RGCs’ axons refine their terminal arbors slowly during postnatal development to attain specific laminar positions in their target areas (Kim et al., 2010; Hong et al., 2014; Osterhout et al., 2014). We find that tSbC RGC axons establish mature lamination patterns shortly after invading their targets (Figs. 5-8). Brn3 transcription factors are important nodes of the gene regulatory networks that govern RGC differentiation and terminal identity, and Brn3c has been suggested to shape axon projection patterns (Wang et al., 2002; Mu and Klein, 2004; Badea et al., 2009). Using cell-type-specific Brn3c deletion and axon mapping, we find no difference in the patterns of tSbC RGC axons developing with and without Brn3c (Fig. 9). The molecular mechanisms that drive tSbC RGC axon growth, keep tSbC RGC axons poised in the optic tract, make them simultaneously invade their targets, and guide them to the correct laminar position, thus, remain unknown. The developmentally stable and selective genetic intersection we identify for tSbC RGC targeting should facilitate future efforts to uncover these mechanisms.

Finally, we find that tSbC RGCs innervate the ipsi- and the contralateral vLGN and dLGN, but only the contralateral SC and NOT (Fig. 3). This could indicate that only a subset of tSbC RGCs innervates SC and NOT or that some tSbC RGC axons bifurcate at the optic chiasm to innervate targets on both sides of the brain. Both options raise interesting developmental and molecular questions and could be distinguished using single-cell labeling approaches in future studies (Hong et al., 2011; Fernandez et al., 2016; Herrera et al., 2019; Mason and Slavi, 2020).

## Acknowledgments

This work was supported by the National Institutes of Health grants EY027411 (D.K.), EY023341 (D.K.), EY026978 (D.K.), the Grace Nelson Lacy Research Fund (D.K.), and an unrestricted grant (Department of Ophthalmology and Visual Sciences) from Research to Prevent Blindness.

## References

Badea TC, Cahill H, Ecker J, Hattar S, Nathans J (2009) Distinct roles of transcription factors brn3a and brn3b in controlling the development, morphology, and function of retinal ganglion cells. Neuron 61:852–864.

Badea TC, Nathans J (2011) Morphologies of mouse retinal ganglion cells expressing transcription factors Brn3a, Brn3b, and Brn3c: analysis of wild type and mutant cells using genetically-directed sparse labeling. Vision Res 51:269–279.

Badea TC, Wang Y, Nathans J (2003) A noninvasive genetic/pharmacologic strategy for visualizing cell morphology and clonal relationships in the mouse. J Neurosci 23:2314–2322.

Badea TC, Williams J, Smallwood P, Shi M, Motajo O, Nathans J (2012) Combinatorial expression of Brn3 transcription factors in somatosensory neurons: genetic and morphologic analysis. J Neurosci 32:995–1007.

Baden T, Berens P, Franke K, Román Rosón M, Bethge M, Euler T (2016) The functional diversity of retinal ganglion cells in the mouse. Nature 529:345–350.

Bae JA, Mu S, Kim JS, Turner NL, Tartavull I, Kemnitz N, Jordan CS, Norton AD, Silversmith WM, Prentki R, Sorek M, David C, Jones DL, Bland D, Sterling ALR, Park J, Briggman KL, Seung HS, Eyewirers (2018) Digital Museum of Retinal Ganglion Cells with Dense Anatomy and Physiology. Cell 173:1293–1306.e19.

Beltramo R, Scanziani M (2019) A collicular visual cortex: Neocortical space for an ancient midbrain visual structure. Science 363:64–69.

Bennett C, Gale SD, Garrett ME, Newton ML, Callaway EM, Murphy GJ, Olsen SR (2019) Higher-Order Thalamic Circuits Channel Parallel Streams of Visual Information in Mice. Neuron Available at: http://www.sciencedirect.com/science/article/pii/S0896627319301199.

Collewijn H (1975) Direction-selective units in the rabbit’s nucleus of the optic tract. Brain Res 100:489–508.

de Monasterio FM (1978) Properties of ganglion cells with atypical receptive-field organization in retina of macaques. J Neurophysiol 41:1435–1449.

Dhande OS, Estevez ME, Quattrochi LE, El-Danaf RN, Nguyen PL, Berson DM, Huberman AD (2013) Genetic dissection of retinal inputs to brainstem nuclei controlling image stabilization. J Neurosci 33:17797–17813.

Dräger UC, Olsen JF (1980) Origins of crossed and uncrossed retinal projections in pigmented and albino mice. J Comp Neurol 191:383–412.

Erskine L, Herrera E (2007) The retinal ganglion cell axon’s journey: insights into molecular mechanisms of axon guidance. Dev Biol 308:1–14.

Fernandez DC, Chang Y-T, Hattar S, Chen S-K (2016) Architecture of retinal projections to the central circadian pacemaker. Proc Natl Acad Sci U S A 113:6047–6052.

Fratzl A, Koltchev AM, Vissers N, Tan YL, Marques-Smith A, Vanessa Stempel A, Branco T, Hofer SB (2021) Flexible inhibitory control of visually evoked defensive behavior by the ventral lateral geniculate nucleus. Neuron 0 Available at: http://www.cell.com/article/S0896627321006577/abstract [Accessed October 5, 2021].

Goetz J, Jessen ZF, Jacobi A, Mani A, Cooler S, Greer D, Kadri S, Segal J, Shekhar K, Sanes J, Schwartz G (2021) Unified classification of mouse retinal ganglion cells using function, morphology, and gene expression. bioRxiv:2021.06.10.447922 Available at: https://www.biorxiv.org/content/10.1101/2021.06.10.447922v1 [Accessed June 13, 2021].

Gollisch T, Meister M (2010) Eye smarter than scientists believed: neural computations in circuits of the retina. Neuron 65:150–164.

Hattar S, Kumar M, Park A, Tong P, Tung J, Yau K-W, Berson DM (2006) Central projections of melanopsin-expressing retinal ganglion cells in the mouse. J Comp Neurol 497:326–349.

Herrera E, Erskine L, Morenilla-Palao C (2019) Guidance of retinal axons in mammals. Semin Cell Dev Biol 85:48–59.

Hong YK, Kim I-J, Sanes JR (2011) Stereotyped axonal arbors of retinal ganglion cell subsets in the mouse superior colliculus. J Comp Neurol 519:1691–1711.

Hong YK, Park S, Litvina EY, Morales J, Sanes JR, Chen C (2014) Refinement of the retinogeniculate synapse by bouton clustering. Neuron 84:332–339.

Huang L, Xi Y, Peng Y, Yang Y, Huang X, Fu Y, Tao Q, Xiao J, Yuan T, An K, Zhao H, Pu M, Xu F, Xue T, Luo M, So K-F, Ren C (2019) A Visual Circuit Related to Habenula Underlies the Antidepressive Effects of Light Therapy. Neuron Available at: http://www.sciencedirect.com/science/article/pii/S0896627319300649.

Huang X, Huang P, Huang L, Hu Z, Liu X, Shen J, Xi Y, Yang Y, Fu Y, Tao Q, Lin S, Xu A, Xu F, Xue T, So K-F, Li H, Ren C (2020) A Visual Circuit Related to the Nucleus Reuniens for the Spatial-Memory-Promoting Effects of Light Treatment. Neuron Available at: http://dx.doi.org/10.1016/j.neuron.2020.10.023.

Huberman AD, Manu M, Koch SM, Susman MW, Lutz AB, Ullian EM, Baccus SA, Barres BA (2008) Architecture and activity-mediated refinement of axonal projections from a mosaic of genetically identified retinal ganglion cells. Neuron 59:425–438.

Huberman AD, Wei W, Elstrott J, Stafford BK, Feller MB, Barres BA (2009) Genetic identification of an On-Off direction-selective retinal ganglion cell subtype reveals a layer-specific subcortical map of posterior motion. Neuron 62:327–334.

Ito S, Feldheim DA, Litke AM (2017) Segregation of visual response properties in the mouse superior colliculus and their modulation during locomotion. J Neurosci Available at: http://dx.doi.org/10.1523/JNEUROSCI.3689-16.2017.

Jacoby J, Schwartz GW (2018) Typology and Circuitry of Suppressed-by-Contrast Retinal Ganglion Cells. Front Cell Neurosci 12:269.

Jacoby J, Zhu Y, DeVries SH, Schwartz GW (2015) An Amacrine Cell Circuit for Signaling Steady Illumination in the Retina. Cell Rep 13:2663–2670.

Johnson KP, Fitzpatrick MJ, Zhao L, Wang B, McCracken S, Williams PR, Kerschensteiner D (2021) Cell-type-specific binocular vision guides predation in mice. Neuron Available at: http://dx.doi.org/10.1016/j.neuron.2021.03.010.

Kato I, Harada K, Hasegawa T, Igarashi T, Koike Y, Kawasaki T (1986) Role of the nucleus of the optic tract in monkeys in relation to optokinetic nystagmus. Brain Res 364:12–22.

Kerschensteiner D, Guido W (2017) Organization of the dorsal lateral geniculate nucleus in the mouse. Vis Neurosci 34:E008.

Kim I-J, Zhang Y, Meister M, Sanes JR (2010) Laminar restriction of retinal ganglion cell dendrites and axons: subtype-specific developmental patterns revealed with transgenic markers. J Neurosci 30:1452–1462.

Kim I-J, Zhang Y, Yamagata M, Meister M, Sanes JR (2008) Molecular identification of a retinal cell type that responds to upward motion. Nature 452:478–482.

Knöferle J, Koch JC, Ostendorf T, Michel U, Planchamp V, Vutova P, Tönges L, Stadelmann C, Brück W, Bähr M, Lingor P (2010) Mechanisms of acute axonal degeneration in the optic nerve in vivo. Proc Natl Acad Sci U S A 107:6064–6069.

Levick WR (1967) Receptive fields and trigger features of ganglion cells in the visual streak of the rabbits retina. J Physiol 188:285–307.

Liang L, Fratzl A, Goldey G, Ramesh RN, Sugden AU, Morgan JL, Chen C, Andermann ML (2018) A Fine-Scale Functional Logic to Convergence from Retina to Thalamus. Cell 173:1343–1355.e24.

Macé É, Montaldo G, Trenholm S, Cowan C, Brignall A, Urban A, Roska B (2018) Whole-Brain Functional Ultrasound Imaging Reveals Brain Modules for Visuomotor Integration. Neuron 100:1241–1251.e7.

Madisen L, Zwingman TA, Sunkin SM, Oh SW, Zariwala HA, Gu H, Ng LL, Palmiter RD, Hawrylycz MJ, Jones AR, Lein ES, Zeng H (2010) A robust and high-throughput Cre reporting and characterization system for the whole mouse brain. Nat Neurosci 13:133–140.

Magnin M, Cooper HM, Mick G (1989) Retinohypothalamic pathway: a breach in the law of Newton-Müller-Gudden? Brain Res 488:390–397.

Martersteck EM, Hirokawa KE, Evarts M, Bernard A, Duan X, Li Y, Ng L, Oh SW, Ouellette B, Royall JJ, Stoecklin M, Wang Q, Zeng H, Sanes JR, Harris JA (2017) Diverse Central Projection Patterns of Retinal Ganglion Cells. Cell Rep 18:2058–2072.

Masland RH, Martin PR (2007) The unsolved mystery of vision. Curr Biol 17:R577–82.

Mason C, Slavi N (2020) Retinal Ganglion Cell Axon Wiring Establishing the Binocular Circuit. Annu Rev Vis Sci 6:215–236.

Moore DL, Goldberg JL (2011) Multiple transcription factor families regulate axon growth and regeneration. Dev Neurobiol 71:1186–1211.

Morin LP, Studholme KM (2014) Retinofugal projections in the mouse. J Comp Neurol 522:3733–3753.

Mu X, Klein WH (2004) A gene regulatory hierarchy for retinal ganglion cell specification and differentiation. Semin Cell Dev Biol 15:115–123.

Niell CM, Stryker MP (2008) Highly selective receptive fields in mouse visual cortex. J Neurosci 28:7520–7536.

Osterhout JA, El-Danaf RN, Nguyen PL, Huberman AD (2014) Birthdate and outgrowth timing predict cellular mechanisms of axon target matching in the developing visual pathway. Cell Rep 8:1006–1017.

Osterhout JA, Josten N, Yamada J, Pan F, Wu S-W, Nguyen PL, Panagiotakos G, Inoue YU, Egusa SF, Volgyi B, Inoue T, Bloomfield SA, Barres BA, Berson DM, Feldheim DA, Huberman AD (2011) Cadherin-6 mediates axon-target matching in a non-image-forming visual circuit. Neuron 71:632–639.

Osterhout JA, Stafford BK, Nguyen PL, Yoshihara Y, Huberman AD (2015) Contactin-4 mediates axon-target specificity and functional development of the accessory optic system. Neuron 86:985–999.

Parmhans N, Fuller AD, Nguyen E, Chuang K, Swygart D, Wienbar SR, Lin T, Kozmik Z, Dong L, Schwartz GW, Badea TC (2021) Identification of retinal ganglion cell types and brain nuclei expressing the transcription factor Brn3c/Pou4f3 using a Cre recombinase knock-in allele. J Comp Neurol 529:1926–1953.

Parmhans N, Sajgo S, Niu J, Luo W, Badea TC (2018) Characterization of retinal ganglion cell, horizontal cell, and amacrine cell types expressing the neurotrophic receptor tyrosine kinase Ret. J Comp Neurol 526:742–766.

Piscopo DM, El-Danaf RN, Huberman AD, Niell CM (2013) Diverse visual features encoded in mouse lateral geniculate nucleus. J Neurosci 33:4642–4656.

Pu ML, Amthor FR (1990) Dendritic morphologies of retinal ganglion cells projecting to the nucleus of the optic tract in the rabbit. J Comp Neurol 302:657–674.

Reese BE (1988) “Hidden lamination” in the dorsal lateral geniculate nucleus: the functional organization of this thalamic region in the rat. Brain Res 472:119–137.

Rheaume BA, Jereen A, Bolisetty M, Sajid MS, Yang Y, Renna K, Sun L, Robson P, Trakhtenberg EF (2018) Single cell transcriptome profiling of retinal ganglion cells identifies cellular subtypes. Nat Commun 9:2759.

Rodieck RW (1967) Receptive fields in the cat retina: a new type. Science 157:90–92.

Rodieck RW (1991) The density recovery profile: A method for the analysis of points in the plane applicable to retinal studies. Visual Neuroscience 6:95–111 Available at: http://dx.doi.org/10.1017/s095252380001049x.

Sabbagh U, Govindaiah G, Somaiya RD, Ha RV, Wei JC, Guido W, Fox MA (2020) Diverse GABAergic neurons organize into subtype-specific sublaminae in the ventral lateral geniculate nucleus. J Neurochem Available at: https://onlinelibrary.wiley.com/doi/abs/10.1111/jnc.15101.

Salay LD, Huberman AD (2021) Divergent outputs of the ventral lateral geniculate nucleus mediate visually evoked defensive behaviors. Cell Rep 37 Available at: http://www.cell.com/article/S2211124721012523/abstract [Accessed October 5, 2021].

Schindelin J, Arganda-Carreras I, Frise E, Kaynig V, Longair M, Pietzsch T, Preibisch S, Rueden C, Saalfeld S, Schmid B, Tinevez J-Y, White DJ, Hartenstein V, Eliceiri K, Tomancak P, Cardona A (2012) Fiji: an open-source platform for biological-image analysis. Nat Methods 9:676–682.

Schwartz G (2021) Retinal Computation. Academic Press.

Simpson JI (1984) THE ACCESSORY OPTIC SYSTEM. Annu Rev Neurosci 7:13–41.

Su J, Sabbagh U, Liang Y, Olejníková L, Dixon KG, Russell AL, Chen J, Pan YA, Triplett JW, Fox MA (2021) A cell-ECM mechanism for connecting the ipsilateral eye to the brain. Proc Natl Acad Sci U S A 118 Available at: http://dx.doi.org/10.1073/pnas.2104343118.

Taniguchi H, He M, Wu P, Kim S, Paik R, Sugino K, Kvitsiani D, Kvitsani D, Fu Y, Lu J, Lin Y, Miyoshi G, Shima Y, Fishell G, Nelson SB, Huang ZJ (2011) A resource of Cre driver lines for genetic targeting of GABAergic neurons in cerebral cortex. Neuron 71:995–1013.

Tien N-W, Pearson JT, Heller CR, Demas J, Kerschensteiner D (2015) Genetically Identified Suppressed-by-Contrast Retinal Ganglion Cells Reliably Signal Self-Generated Visual Stimuli. J Neurosci 35:10815–10820.

Tran NM, Shekhar K, Whitney IE, Jacobi A, Benhar I, Hong G, Yan W, Adiconis X, Arnold ME, Lee JM, Levin JZ, Lin D, Wang C, Lieber CM, Regev A, He Z, Sanes JR (2019) Single-Cell Profiles of Retinal Ganglion Cells Differing in Resilience to Injury Reveal Neuroprotective Genes. Neuron 104:1039–1055.e12.

Wang SW, Mu X, Bowers WJ, Kim D-S, Plas DJ, Crair MC, Federoff HJ, Gan L, Klein WH (2002) Brn3b/Brn3c double knockout mice reveal an unsuspected role for Brn3c in retinal ganglion cell axon outgrowth. Development 129:467–477.

Williams PR, Benowitz LI, Goldberg JL, He Z (2020) Axon Regeneration in the Mammalian Optic Nerve. Annu Rev Vis Sci 6:195–213.

Xiang M, Zhou L, Macke JP, Yoshioka T, Hendry SH, Eddy RL, Shows TB, Nathans J (1995) The Brn-3 family of POU-domain factors: primary structure, binding specificity, and expression in subsets of retinal ganglion cells and somatosensory neurons. J Neurosci 15:4762–4785.

Yakushin SB, Gizzi M, Reisine H, Raphan T, Büttner-Ennever J, Cohen B (2000) Functions of the nucleus of the optic tract (NOT). II. Control of ocular pursuit. Exp Brain Res 131:433–447.

Yonehara K, Shintani T, Suzuki R, Sakuta H, Takeuchi Y, Nakamura-Yonehara K, Noda M (2008) Expression of SPIG1 reveals development of a retinal ganglion cell subtype projecting to the medial terminal nucleus in the mouse. PLoS One 3:e1533.

